# NASC-seq monitors RNA synthesis in single cells

**DOI:** 10.1101/498667

**Authors:** Gert-Jan Hendriks, Lisa A. Jung, Anton J.M. Larsson, Oscar Andersson Forsman, Michael Lidschreiber, Katja Lidschreiber, Patrick Cramer, Rickard Sandberg

**Affiliations:** Karolinska Institutet, Department of Cell and Molecular Biology, Biomedicum, Solnavägen 9, 171 65 Solna, Stockholm, Sweden; Karolinska Institutet, Department of Biosciences and Nutrition and Center for Innovative Medicine and Science for Life Laboratory, NEO, Hälsovägen 7C, 141 83 Huddinge, Sweden; Department of Molecular Biology, Max Planck Institute for Biophysical Chemistry, Am Fassberg 11, 37077 Göttingen, Germany

## Abstract

Sequencing of newly synthesized RNA can monitor transcriptional dynamics with great sensitivity and high temporal resolution, but is currently restricted to populations of cells. Here, we developed <u>n</u>ewly synthesized <u>a</u>lkylation-dependent <u>s</u>ingle-<u>c</u>ell RNA sequencing (NASC-seq), to monitor both newly synthesized and pre-existing RNA in single cells. We validated the method on pre-alkylated exogenous spike-in RNA, and by demonstrating that more newly synthesized RNA was detected for genes with known high mRNA turnover. Importantly, NASC-seq reveals rapidly up- and down-regulated genes during the T-cell activation, and RNA sequenced for induced genes were essentially only newly synthesized. Moreover, the newly synthesized and preexisting transcriptomes after T-cell activation were distinct confirming that we indeed could simultaneously measure gene expression corresponding to two time points in single cells. Altogether, NASC-seq is a powerful tool to investigate transcriptional dynamics and it will enable the precise monitoring of RNA synthesis at flexible time periods during homeostasis, perturbation responses and cellular differentiation.

## Introduction

Whereas RNA-seq measures cellular RNA levels, insights into the kinetics of RNA transcription, processing, and degradation rely on metabolic labeling and sequencing of newly synthesized RNA^1–3^. RNA labelling methods enable the detection of rapid transcriptional responses to cellular stimuli or perturbations^4, 5^. They generally rely on the incorporation of 4-thiouridine (4sU) into newly synthesized RNA during gene transcription and subsequent biochemical separation of 4sU-labeled and unlabeled RNA. As these methods require large amounts of total cellular RNA they have been limited to studies of average cellular responses across cell populations. Recent advances in single-cell RNA sequencing and data analysis have however revealed that responses to stimuli are not uniform across cells^6^.

Thus a method is required to monitor transcriptional dynamics in single cells by sequencing of newly synthesized RNA. Although 4sU-labeled RNA cannot be isolated from single cells in quantities that allow for its sequencing, recent cell population experiments showed that the need to separate labeled and unlabeled RNA can be eliminated. The recent approaches rely on chemical modification of 4sU residues in total cellular RNA, which leads to apparent T-C transitions during reverse transcription that can be read out by sequencing^7–10^. Newly synthesized RNA thus can then be separated from pre-existing RNA *in silico* during computational analysis of the sequencing data.

## Results

Based on these findings we sought to combine 4sU labeling, chemical conversion of the 4sU residue, and single-cell RNA-seq. After several rounds of experimental and computational adaptations and optimization, we arrived at a procedure for the sequencing of newly synthesized and pre-existing RNA in single cells that we termed NASC-seq, for new transcriptome alkylation-dependent single-cell sequencing (**Fig. 1a**). Briefly, cells are first exposed to 4sU, and then sorted and lysed individually. RNA is immobilized on magnetic beads with a biotinylated oligo-dT primer^11^ and is then alkylated. Single-cell RNA-seq libraries are then constructed using a modified version of Smart-seq2^12^. Reverse transcription over alkylated 4sU residues triggers the misincorporation of guanines instead of adenosines, leading to T-C transitions that identify newly synthesized RNA in the sequenced libraries.

**Figure 1.**
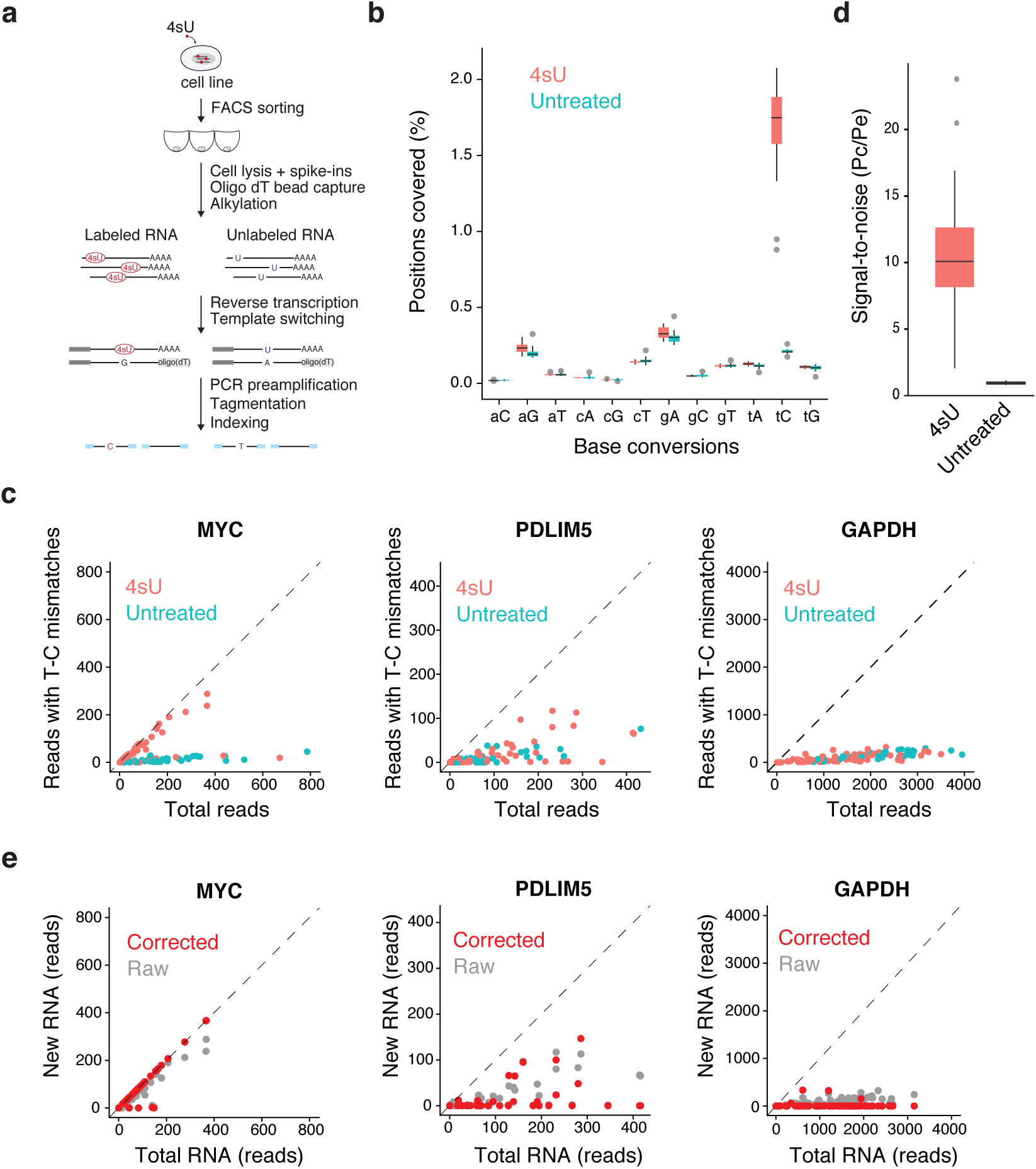
Global sequencing of newly synthesized RNA in single cells. **(a)** Illustration of the NASC-seq methodology. In brief, alkylation is performed on RNAs immobilized on beads for the subsequent wash before proceeding with standard PCR, tagmentation and construction of sequencing libraries. **(b)** Observed conversion rates in K562 cells treated with 4sU (50μM, 3 hours; red, 16 cells) or untreated (blue, 14 cells) on the positive strand within genes. T-C (tC) conversions are significantly (P= 1.375e-08, Mann-Whitney *U*-test, two-sided) increased in cells treated with 4sU. The line in the boxplot indicates the median value, the two hinges display the first and third quartiles. The whiskers range from the hinges to the highest or lowest point that is no further than 1.5 x the interquartile range. **(c)** Scatter plots showing the total number of sequenced reads (x-axis) against the number of reads with T-C conversions (y-axis) for the MYC, PDLIM5 and GAPDH genes in 4sU treated (50μM, 1 hr) and untreated cells. **(d)** Signal to noise estimated as the probability of conversion divided by the probability of error, for K562 cells exposed to 50uM for 1 hour (n=75) compared to K562 cells that were not exposed to 4sU (n=41). Median, hinges and whiskers are shown as in **(b)**.**(e)** Scatter plots showing newly synthesized (new) RNA inferred using the mixture-model against total RNA (i.e. number of reads) for the same genes and 4sU treated cells from **(c)** that passed the post-correction QC filter (see methods).

We first applied NASC-seq to human K562 cells. We exposed cells for 180 minutes to increasing concentrations of 4sU and constructed sequencing libraries. The dominating type of base conversion that we observed was the T-C transition (**Fig. 1b**), demonstrating that a strong signal was present in the data (**Fig. 1b**). Based on the observed transitions, the frequency of 4sU incorporation at 50 μM 4sU was ~2% and thus comparable to that obtained in cell populations^7^ (**Fig. 1b**). We found that low-input alkylation was equally efficient as bulk alkylation (**Supplemental Fig. 1a**), that 4sU-labeled control (spike-in) RNA contained high levels of T-C transitions (**Supplemental Fig. 1b**), and that higher concentrations of 4sU lowered the transcriptome complexity in libraries, observed as lowered numbers of genes detected per cell (**Supplemental Fig. 1c**). After optimizing the experimental parameters, we arrived at a robust protocol for NASC-seq.

To further investigate whether NASC-seq reveals newly synthesized RNA, we analyzed genes that encode mRNAs with high, intermediate, and low turnover as determined in previous population measurements^3^. We reasoned that mRNAs with high turnover have a larger fraction of newly synthesized RNA and this should be indicated by a higher number of T-C transitions in NASC-seq data. Indeed, comparing the numbers of reads with T-C transitions in genes with high (MYC), intermediate (PDLIM5) or low (GAPDH) RNA turnover revealed that the detected RNAs had high, intermediary, and low numbers of T-C transitions, respectively (**Fig. 1c; Supplemental Fig. 1d**).

This analysis indicated that the number of reads with T-C transitions is a proxy for the amount of newly synthesized RNA. However, a fraction of these reads are apparently false positives because 5–10% of the total reads from unlabeled cells showed T-C transitions that are likely caused by sequencing and PCR errors (**Fig. 1b**). To reduce the number of false positive reads, we adapted a binomial mixture model from the recently published GRAND-SLAM statistical approach^13^. The approach also describes an estimation of true conversion probability (*p_c_*) based on the background error probability (*p_e_*). This approach improved the detection of newly synthesized RNA (**Fig. 1c,e**) and revealed that we obtained a signal-to-noise ratio of ~10 in this experiment (**Fig. 1d**).

We next sorted all genes by their mRNA turnover^3^ and generated groups of genes encoding for mRNAs with low (bottom 20%) and high turnover (top 20%). Genes with high mRNA turnover showed on average more reads with T-C transitions (**Supplemental Fig. 1e**). This global trend was also consistent with NASC-seq detecting newly synthesized RNA.

We now tested whether NASC-seq can monitor changes in newly synthesized RNA in single cells during a transcription response. We labeled Jurkat T-cells with 4sU, and induced a rapid transcriptional response by simultaneous addition of PMA and ionomycin for 30 minutes as described^4^ (**Fig. 2a**). NASC-seq revealed a high number of T-C transitions for genes that were known to be rapidly induced upon stimulation, such as EGR1 and FOS, but not for the non-induced genes GAPDH and ACTB, providing a negative control (**Fig. 2b**). Application of the mixture model essentially eliminated the number of false positive reads with T-C transitions (**Fig. 2b**).

**Figure 2.**
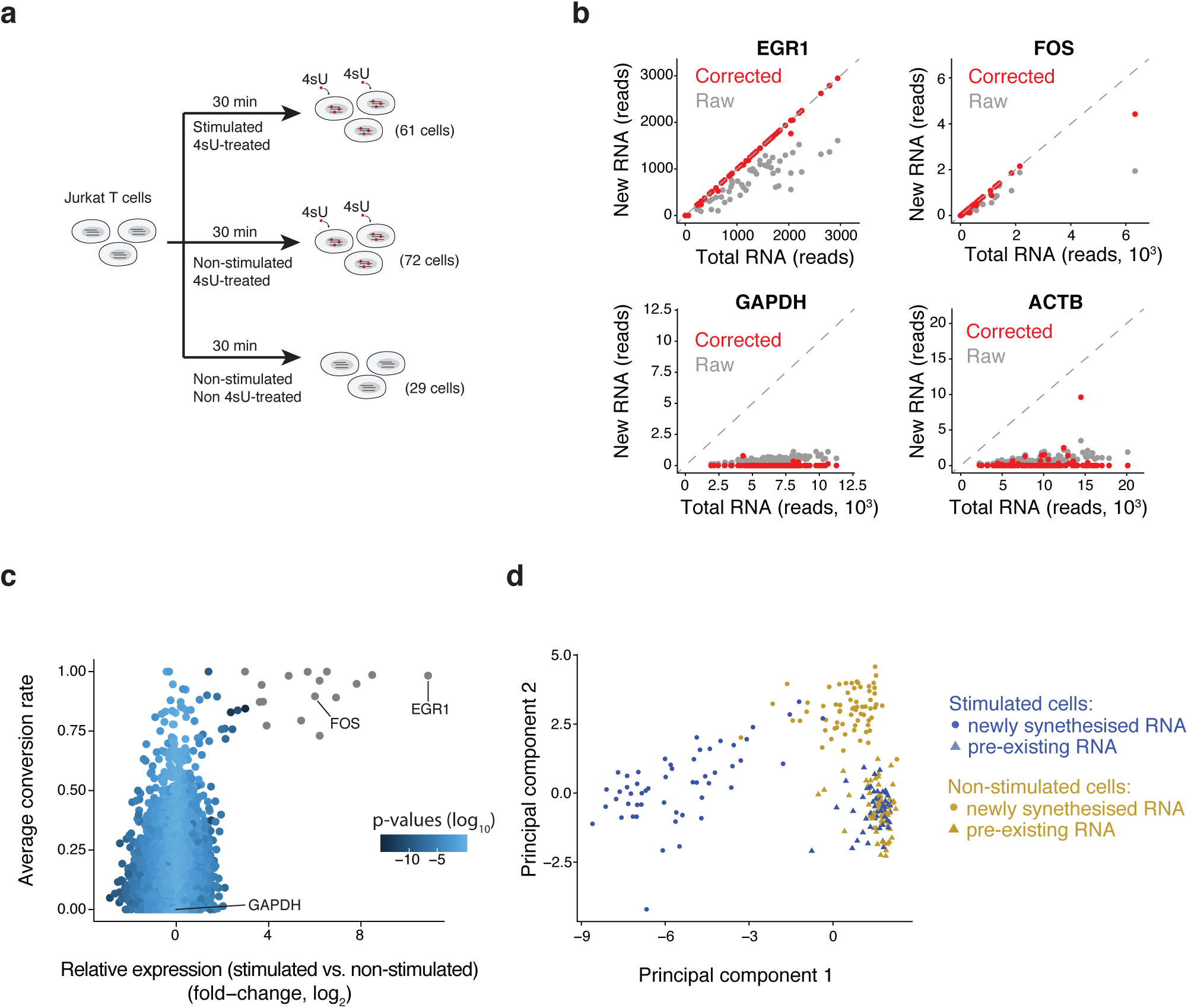
Single-cell analysis of RNA dynamics during T-cell activation. **(a)** Illustration of experiment design. Jurkat T-cells stimulated with PMA and ionomycin where simultaneously exposed to 4sU for 30 minutes. Unstimulated cells received only 4sU and no PMA and ionomycin. Untreated cells were collected that were neither stimulated nor labeled with 4sU. **(b)** Scatter plots of total and new RNA for two response genes (EGR1 and FOS; positive controls) and two lowly turned-over genes (GAPDH and ACTB) for all 4sU treated cells (total n=119, PMA/ionomycin stimulated n = 53, unstimulated n =66). **(c)** Scatter plots showing differential expression of genes with more than 5 *π_g_* estimates (n=8966) in Jurkat cells stimulated with PMA and ionomycin, plotted against mean conversion rate per cell. EGR1, FOS and GAPDH are pointed out. Genes were colored according differential expression (stimulated vs. nonstimulated cells using ROTS), genes in gray had uncorrected p-values below 10^−15^. **(d)** Two-dimensional PCA plot showing the cellular transcriptomes after separating each cell into newly synthesized and pre-existing RNA for PMA/ionomycin stimulated (n=53) and non-stimulated Jurkat cells (n=66).

Overexpressed genes in stimulated Jurkat cells, compared to non-stimulated cells, generally show high average conversion rates, illustrating the sensitivity of NASC-seq to detect changes in expression kinetics (**Fig. 2c**). To investigate whether NASC-seq could generally detect transcriptionally induced genes upon T-cell stimulation, we selected genes that were significantly up- and down-regulated, respectively, under the same conditions in bulk TT-seq measurements and that were not detected by standard RNA-seq. Indeed, these groups of up- and down-regulated genes showed a significant increase and decrease, respectively, in their NASC-seq signal (**Supplemental Fig. 2b**). Taken together, NASC-seq clearly detected transcriptionally active T-cell genes that are known to be induced under stimulating conditions^14^.

Finally, we sought to analyze the temporal resolution of NASC-seq. We subjected T-cell activation NASC-seq data to principal component analysis (PCA). PCA could indeed separate newly synthesized RNAs in stimulated cells from those in nonstimulated cells (**Fig. 2d**, principal component 1, PC1). This separation was much less pronounced when total RNA measurements were used (**Supplemental Fig. 2a**). Also, pre-existing RNAs did not separate in this analysis, as expected (**Fig. 2d**). Together these analyses show that NASC-seq can effectively measure the transcriptome at two time points per cell and it is therefore very well suited to monitor rapid changes in transcription activity in single cells.

## Discussion

In this study, we introduce a novel method, NASC-seq, that monitors newly synthesized RNA in single human cells. NASC-seq is based on a combination of RNA labeling with 4sU, RNA modification by alkylation as in SLAM-seq^7^, RNA sequencing library preparation as in Smart-seq2, and data analysis that includes a computational model from GRAND-SLAM^13^. We validate NASC-seq by comparison with RNA labeling data in cell populations. We show that NASC-seq can separate newly synthesized from pre-existing RNA in single human cells, and that it can also monitor up- and down-regulation of transcription during a rapid cellular response.

Recent studies showed that information on RNA synthesis can be obtained from the detection of intronic sequence in single cells^15^, ^16^ and can inform on future cell fate^15^. NASC-seq expands on these possibilities by measuring newly synthesized RNA from periods of 4sU exposure without being restricted to genes for which intronic RNA reads are detected or limited in time by the kinetics of pre-mRNA splicing. We note that measuring transcriptomes at two time points per cell could provide new insights into transcriptional kinetics^17^. Thus NASC-seq is ideally suited to monitor changes in transcription activity with high sensitivity and temporal resolution during cell differentiation, tissue engineering, and organism development.

## Author contributions

GJH and LAJ established and carried out NASC-seq experiments; KL, GJH and RS designed the NASC-seq approach; GJH, AL, OAF and ML developed computational methodology; AL and OAF implemented the statistical inference approach; GJH, AL, ML and LAJ analysed data; LAJ carried out TT-seq experiments; GJH, LAJ, PC and RS wrote the manuscript with input from all authors; PC and RS supervised the study.

## Acknowledgements

We are grateful to Christoph Ziegenhain and the other members of the Sandberg laboratory for fruitful discussions. We thank Saskia Gressel, Kerstin Maier and Noah Mottelson (MPI-BPC Göttingen) for help. GJH was funded by EMBO long-term fellowship ALTF 1528–2016 and HFSP long-term fellowship LT000155/2017-L. PC was funded by the Center for Innovative Medicine (CIMED), the Advanced Grant TRANSREGULON from the European Research Council, and the Volkswagen Foundation. RS was funded by the European Research Council (648842), the Swedish Research Council (2017–01062), the Knut and Alice Wallenberg’s foundation (2017.0110) and the Bert L. and N. Kuggie Vallee Foundation.

## Accession codes

The sequencing data and processed files were deposited in the GEO database under accession codes (id pending).

## Code availability

The statistical model and code for inference of newly synthesized and pre-existing RNA will be provided at Github (https://github.com/sandberg-lab/NASC-seq).

## Supplemental Figure Legends

**Figure S1.**
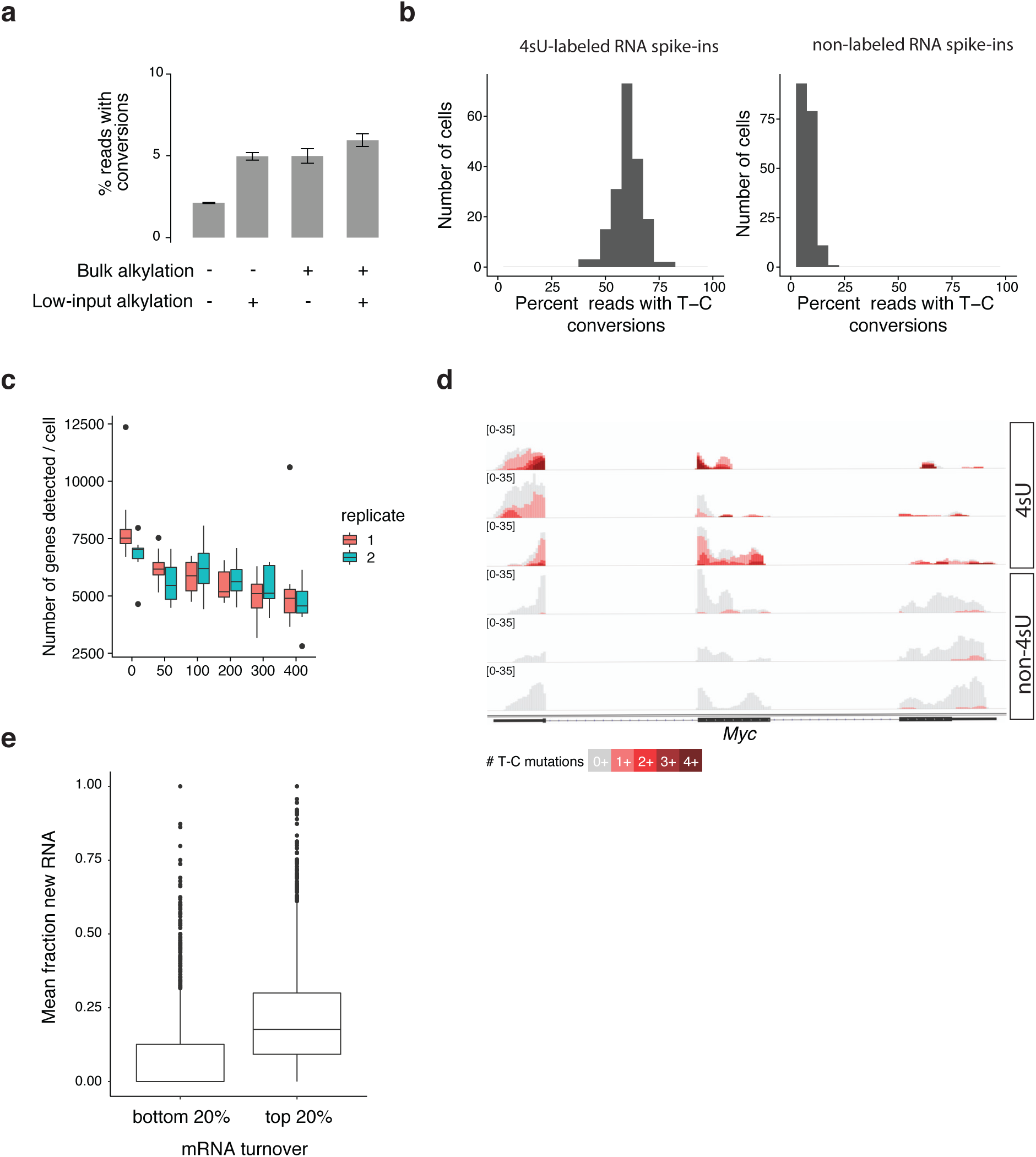
Detailed characterization of NASC-seq on K562 cells. **(A)** Comparison of bulk and low-input alkylation. 50 pg bulk RNA from K562 cells and single K562 cells, both treated for 3h with 500 uM 4sU, were subjected to alkylation either during the NASC-seq protocol (low-input) or previously in bulk format. Shown is the percentage (%) of reads with conversions. Error bars indicate standard deviation. **(B)** Verification of T-C transitions in 4sU-labeld control RNA. Spike-in RNA was generated either with (4sU spike-in RNA) or without 4sU addition (non-4sU spike-in RNA). Shown are % T-C conversions. **(C)** Numbers of genes detected per cell for total single-cell RNA-seq data from K562 cells labeled with increasing doses of 4sU for 180 minutes labeling time. At least one read has to map to a gene for it to be counted as detected. The colors indicate two biological replicates. **(D)** Representative genome browser tracks of the *MYC* gene of three 4sU-treated and three untreated K562 cells showing coverage from all reads (grey) and coverage from reads with at least one or more T-C mutations (red, darker color indicates higher number of T-C mutations, see legend). **(E)** Distribution of fraction of new RNA for genes with low (bottom 20%, n = 2099, left) and high (top 20%, n = 2100, right) turnover rates. Turnover rate was defined as synthesis rate * decay rate using rates estimated from bulk TT-seq data^3^. Only genes with a total read sum over all cells of at least 100 in NASC-seq data were considered. The P-value (<2.2e-16) was derived by two-sided Mann–Whitney *U*-test. Box limits are the first and third quartiles, the band inside the box is the median. The ends of the whiskers extend the box by 1.5 times the interquartile range.

**Figure S2.**
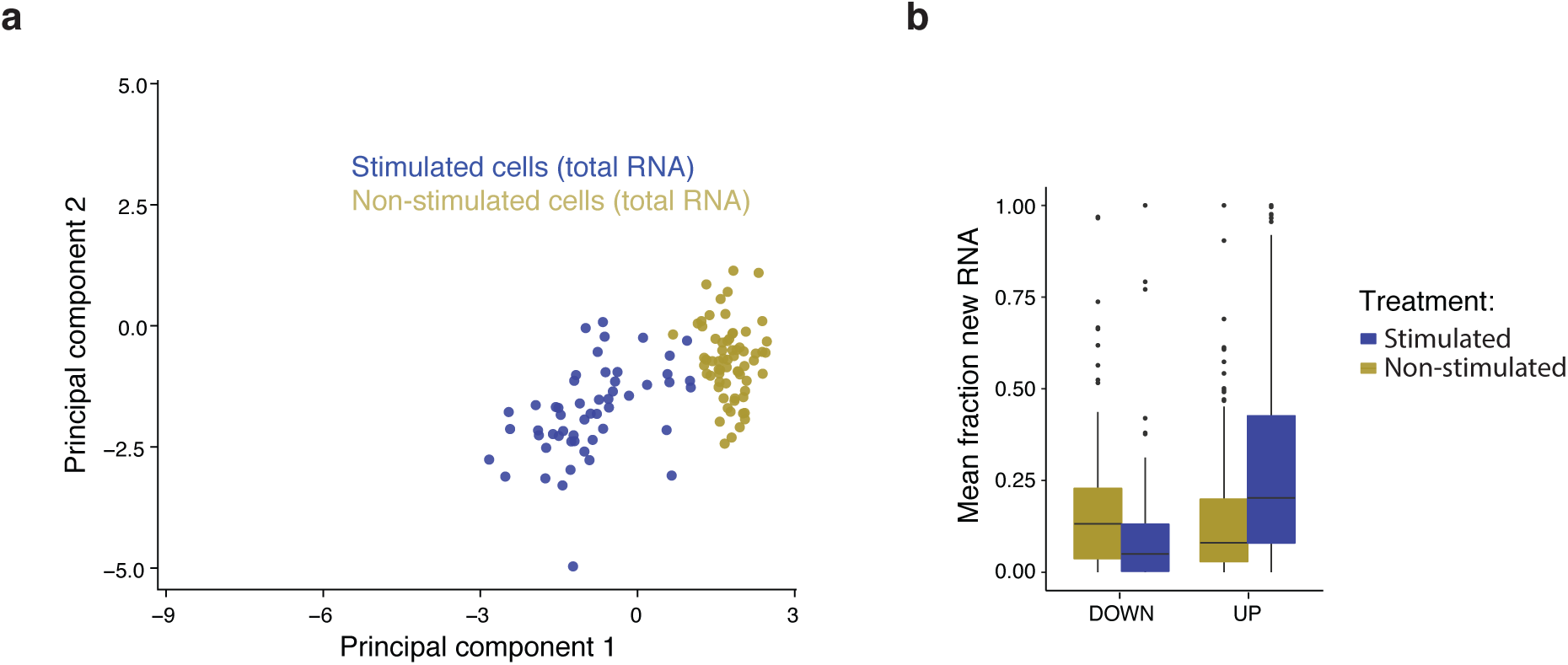
Extended NASC-seq analysis of T-cell activation. **(A)** Principal component analysis (PCA) for total RNA from Jurkat cells stimulated with PMA and ionomycin, and unstimulated cells. Axes are scaled to match axes in figure 2D. **(B)** Distribution of mean fraction of new RNA for genes which are detected as significantly up-regulated (right, n = 116) and down-regulated (left, n = 189) upon T-cell stimulation (for 30 min with PMA and ionomycin) in newly synthesized RNA but not total RNA taken from a TT-seq experiment (see Methods). P-values (lower= 5.299e-07, upper= 2.959e-14) were derived by two-sided Mann–Whitney *U*-test. Box limits are the first and third quartiles, the band inside the box is the median. The ends of the whiskers extend the box by 1.5 times the interquartile range.

## Methods

### Generation of poly-A RNA spike-ins

RNA spike-ins (ERCC-00043, ERCC-00170, ERCC-00136, ERCC-00145, ERCC-00092 and ERCC-00002) were generated as described previously^3^, except reverse transcription was done from plasmid DNA also encoding a poly-A-tail.

### Cell culture and stimulation

Cells were grown in RPMI 1640 medium (Gibco) supplemented with 10% fetal bovine serum (Sigma) and 1% Penicillin/Streptomycin (HyClone) at 37°C under 5% CO_2_. Jurkat cells (E6.1 clone) were acquired from ATCC, K562 cells from DSMZ (Braunschweig, Germany). Cells were routinely tested for mycoplasma contamination (MycoAlert, Lonza). On the day of the experiment, Jurkat cells were seeded to a density of 1*10^6^ cells/ml and stimulated with 50ng/μl phorbol-12-myristate-13-acetate (PMA) and 1μM ionomycin (Sigma) for 30 min at 37°C under 5% CO_2_.

### Bulk RNA alkylation

K562 cells were treated with 500μM 4-thiouridine (4sU, Sigma) at 37°C under 5% CO_2_ for 3 h. Total RNA was isolated using TRIzol (Life Technologies) according to the manufacturer’s instructions. RNA spike-in mix was added during RNA isolation. For bulk alkylation, 5μg of RNA was alkylated as described in Herzog et al.^7^ and purified using isopropanol precipitation. RNA was analyzed using a Bioanalyzer 2100 system (Agilent). Low-input alkylation was performed as described below (NASC-seq).

### TT-seq

Two biological replicates of TT-seq reactions were performed as described previously^3^. Jurkat cells were treated for 30 min with solvent control (DMSO, Sigma) or with PMA and ionomycin. During the last 5 minutes of each time point cells were labelled in media with 500μM 4-thiouridine (4sU, Sigma) at 37°C under 5% CO_2_. Cells were harvested by centrifugation at 1400xg for 2 min. Total RNA was extracted using TRIzol (Life Technologies) according to the manufacturer’s instructions under the addition of spike-ins. RNAs were sonicated using in a Bioruptor Plus instrument (Diagenode). 4sU-labeled RNA was purified from 300μg total fragmented RNA. Separation of labeled RNA was achieved with streptavidin beads (Miltenyi Biotec). Prior to library preparation, 4sU-labeled RNA treated with DNase (Qiagen), purified (miRNeasy Micro Kit, Qiagen) and quantified. Strand-specific libraries were prepared with the Ovation Universal RNA-Seq System (NuGEN). The size-selected and pre-amplified fragments were analyzed on a Bioanalyzer 2100 (Agilent). Samples were sequenced on an Illumina NextSeq 550 instrument. Data analysis was performed essentially as in Michel et al.^4^. Briefly, paired-end 75 bp reads were mapped with STAR^18^ (version 2.6.0c) to the hg38 (GRCh38) genome assembly (Human Genome Reference Consortium). Gene expression fold changes upon T-cell stimulation for each time point were calculated using the R/Bioconductor implementation of DESeq2^19^. Differentially expressed genes were identified applying a fold change cutoff of 2 and an adjusted P-value cutoff of 0.05 comparing sample to solvent control measurements. For comparison of NASC-seq data with TT-seq data, we calculated the fraction of new RNA (Figures S1e and S2b) by taking the sum of reads from newly synthesized RNA divided by the sum of total reads over all cells, thereby creating an *in silico* bulk for better comparison with bulk TT-seq data. K562 TT-seq data was taken from Schwalb et al.^3^ (GSE75792).

### NASC-seq

Cells were labeled in medium with 4sU (Sigma), washed with cold PBS and sorted into lysis buffer (3ul, 166mM Sodium Phosphate pH 8.0, RNase inhibitor, spike-in RNAs) in wells of PCR plates. Plates were frozen at −80°C until use. Streptavidin beads (MyOne Dynabeads Strepavidin C1) were washed once with buffer 1 (0.1M NaOH, 0.05M NaCl), twice with buffer 2 (0.1M NaOH), and once with 2x B&W buffer (2M NaCl, 10mM Tris-HCl pH7.4, 1mM EDTA) before the binding reaction (1x B&W, 50uM oligo-dT). Beads were incubated with agitation at room temperature for 15 minutes and washed twice with 1x B&W. Cells were lysed at 80°C for 3 minutes and beads were added to the cells in a volume of 2ul. RNA was bound during 20 minutes of incubation at room temperature on a thermoshaker (Eppendorf ThermoMixer C). 5ul of alkylation mix (20mM IAA in DMSO) was added for a final alkylation reaction of 50mM Sodium Phosphate pH 8.0, 10mM IAA, 50% DMSO. Alkylation was stopped by adding STOP solution (2x Superscript II buffer,0.3% Tween 20, 60mM DTT), incubating for 5 minutes on a magnet, and removing the supernatant from the beads containing the alkylated RNA. RT and the remaining library preparation was performed according to a modified version of Smart-seq2^18^. The modifications included the removal of the inactivation step from the RT thermocycling program, performing the RT on a thermoshaker (Eppendorf ThermoMixer C) and the use of custom barcoded primers for the Nextera tagmentation. The resulting libraries were sequenced on a NextSeq500 instrument (Illumina) using either single-end (75-cycle) or paired-end (300-cycle) sequencing strategies.

### Computational pipeline

After sequencing the resulting bcl files were demultiplexed to fastq files with *bcl2fastq* (Illumina). Then we trimmed the nextera adapters with *TrimGalore^20^* (v 0.4.5). The trimmed fastq files were then aligned to the hg38 human genome using STAR^18^ (v 2.5). We removed duplicates using the MarkDuplicates command in *Picard* (v 2.17.6). We then annotated the gene each reads maps to using *FeatureCounts^21^* (v 1.6.2). We further annotated all mismatches to the reference genome within each read with the location of the mismatch for the T-C or A-G mismatches depending on the strand of the annotated gene (plus and negative respectively). We then considered mismatches in positions which appear in a high frequency over many cells and marked those positions as single nucleotide variants (SNV) to be ignored. We then re-annotated the mismatches for each read with the SNV positions ignored and avoided double counting in overlapping paired end reads. Low quality cells were filtered out depending according to the following requirements; for Jurkat cell experiments, each cell was required to have 1,000,000 reads, with at least 50% mapping to features. For K562 experiments, each cell was required to have 800,000 reads, and a minimal of 500,000 and 30% mapping to features.

We then estimated the probability of a given position being converted in a new read (*p_c_*) based on the background probability of a mismatch due to error (*p_e_*) for each cell. We estimated *p_e_* by calculating the mean fraction of C-T and G-A mismatches in the given cell, since we concluded that these mismatches follow agree with the T-C and A-G mismatches in untreated cells. To estimate *p_c_* we implemented the Expectation-Maximization algorithm described in Jürges et al^13^. For each gene in each cell, we estimated the proportion of new reads, *π_g_* using the binomial mixture model described in GRAND-SLAM^13^ (see below). We only ran the estimation if the gene had a minimum of 16 reads mapped to ensure reliable estimates. If the gene had more than 1000 mapped reads, we subsampled down to 1000 reads to shorten runtime.

### Estimating the proportion of reads originating from newly transcribed transcripts

To estimate the proportion of new reads for each gene, *π_g_*, we implemented the binomial mixture model described in Jürges et al^13^. In the mixture model, each mismatch is either due to a conversion with probability *p_c_* or an error with probability *p_e_*. The probability of *y* positions having mismatches in a read containing *n* positions which may be converted is

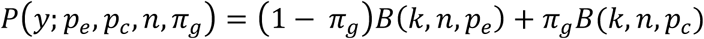

where *B*(*k, n, p*) is the binomial probability mass function.

We estimated *π_g_* by building a generative model in the STAN modelling language.

We use a beta prior for *π_g_* with hyperparameters *α* and *β*

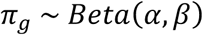

and estimate *π_g_* by maximizing the log-likelihood

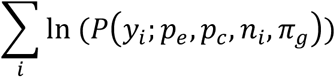

where each index *i* indicates a read for that gene. The hyperparameters were log-transformed and both initialized at 0, while *π_g_* was initialized at 0.5. We will then estimate the mean of the beta distribution, which cannot be 0 or 1 by definition. The mode is therefore more appropriate, which we can calculate by

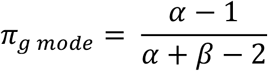

for *α,β >* 1. The mode is 1 if *α >* 1, *β* < 1 and 0 if *α <* 1, *β* > 1. The other possible cases do not occur in our estimation procedure.

Cells with an average standard deviation of the *π_g mode_* that was higher than 0.1 were filtered out and not used for downstream analysis.

